# Computational tool to study perturbations in muscle regulation and its application to heart disease

**DOI:** 10.1101/548404

**Authors:** Samantha K. Barrick, Sarah R. Clippinger, Lina Greenberg, Michael J. Greenberg

## Abstract

Striated muscle contraction occurs when myosin thick filaments bind to thin filaments in the sarcomere and generate pulling forces. This process is regulated by calcium, and it can be perturbed by pathological conditions (e.g., myopathies), physiological adaptations (e.g., β-adrenergic stimulation), and pharmacological interventions. Therefore, it is important to have a methodology to robustly determine the mechanism of these perturbations and statistically evaluate their effects. Here, we present an approach to measure the equilibrium constants that govern muscle activation, estimate uncertainty in these parameters, and statistically test the effects of perturbations. We provide a MATLAB-based computational tool for these analyses, along with easy-to-follow tutorials that make this approach accessible. The hypothesis testing and error estimation approaches described here are broadly applicable, and the provided tools work with other types of data, including cellular measurements. To demonstrate the utility of the approach, we apply it to determine the biophysical mechanism of a mutation that causes familial hypertrophic cardiomyopathy. This approach is generally useful for studying the mechanisms of muscle diseases and therapeutic interventions that target muscle contraction.

## Introduction

Force production in cardiac and skeletal muscle is tightly regulated to ensure that contraction occurs in a controlled and concerted manner. Dysfunction of this regulation can lead to a wide array of diseases, including cardiomyopathies, and there are currently several therapies in development that target this regulation (1–3). Given the role of perturbations of muscle regulation in health and disease, there is an outstanding need for tools that can resolve statistically significant changes in this regulation.

At the molecular scale, force production in muscle is powered by the molecular motor myosin, which contracts the sarcomere by pulling thin filaments (i.e., actin filaments decorated with tropomyosin and the troponin complex) toward the M-line of the sarcomere. The interaction between myosin and the thin filament is regulated in a calcium-dependent manner, where calcium influx into the cytoplasm leads to activation of the thin filament and subsequent muscle contraction. In their landmark work, McKillop and Geeves (4) used a battery of biochemical and biophysical techniques to demonstrate that thin-filament activation is a multi-step process, requiring contributions from calcium binding to troponin as well as actomyosin binding. Their model is known as the “three-state model” (Fig. 1). In the absence of calcium, tropomyosin is in the blocked state, obscuring the myosin-binding site on the thin filament and inhibiting muscle contraction. Calcium binding to the troponin complex causes tropomyosin to shift to the closed state on the thin filament, exposing part of the myosin-binding site. Tropomyosin can either move to the open position spontaneously, where it permits myosin strong binding, or it can be pushed there by myosin binding. Myosin first binds weakly to the thin filament, then isomerizes to a strongly bound, force-generating state. The binding of one myosin to the thin filament pushes tropomyosin into the open position, exposing adjacent myosin-binding sites and leading to cooperative activation. The three states of tropomyosin positioning along the thin filament were subsequently confirmed using structural techniques (5).

**Figure 1.**
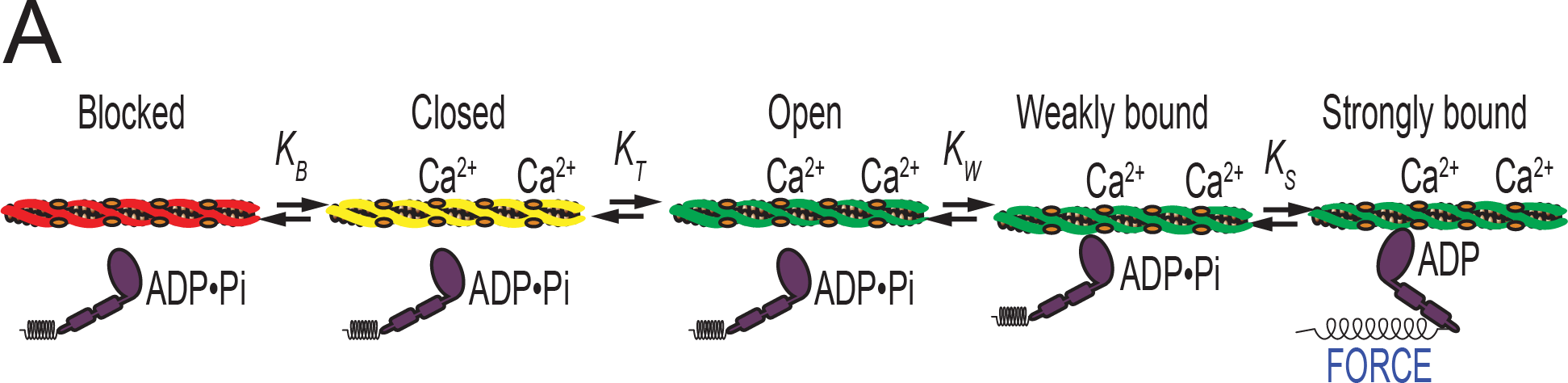
Schematic of the three-state model of muscle regulation. Red, yellow, and green represent the blocked, closed, and open states of tropomyosin, respectively. The equilibrium constants *K*_*B*_ and *K*_*T*_ describe transitions between states of tropomyosin on the thin filament, whereas *K*_*W*_ and *K*_*S*_ describe weak and strong myosin binding, respectively.

The three-state model provides a useful framework to understand the mechanism of perturbations in skeletal and cardiac thin-filament regulation, such as drug treatments (1, 6), protein isoform changes (7), disease-causing mutations (8), and post-translational modifications (9, 10). The formalism laid out by McKillop and Geeves enables the determination of the equilibrium constants that govern transitions between states, and thus the population of thin-filament regulatory units in each state (Fig. 1). However, it has been challenging to precisely define the values of these equilibrium constants, due in part to the number of free parameters in the model. Furthermore, the original McKillop and Geeves approach did not provide a methodology for assessment of uncertainty in parameter values or hypothesis testing. Such a methodology is necessary for the rigorous assessment of the effects of perturbations.

Here, we have modified the approach of McKillop and Geeves to better resolve the effects of perturbations in tropomyosin positioning along the thin filament. We provide a MATLAB-based computational tool and a user guide that enables users to measure the equilibrium constants that govern thin-filament activation, estimate uncertainty in these parameters, and statistically test the effects of perturbations. We demonstrate the utility of our approach by applying it to determine the biophysical mechanism of a mutation in troponin T, ΔE160, that causes familial hypertrophic cardiomyopathy (HCM) (11). This approach allows for the robust determination of differences between wild-type and mutant proteins, which is an important step toward developing novel therapies to treat HCM and other devastating muscle diseases.

## Materials and Methods

### Determination of equilibrium constants using the McKillop and Geeves analysis

In the classic McKillop and Geeves analysis, the values of the equilibrium constants that govern muscle activation (Fig. 1) were determined from biochemical measurements. *K*_B_ was calculated based on stopped-flow measurement of the rate of myosin binding to regulated thin filaments (RTFs) (see Supporting Materials for details). *K_W_*, *K_T_*, and *nH* (i.e., the size of the cooperative unit) were calculated from titrations of pyrene-labeled RTFs with myosin performed at saturating calcium (pCa 3) and in the absence of calcium (2 mM EGTA) (see Supporting Materials for details). The relationship between the fraction of myosin-bound subunits, *f*([m]), and the fractional change in pyrene fluorescence upon myosin binding is given by the following equation (4):

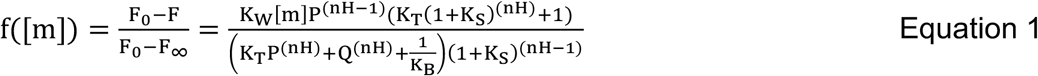

where *F* is the measured pyrene fluorescence; *F*_0_ and F_∞_ are the pyrene fluorescence in the absence of myosin and at saturating myosin, respectively; [m] is the concentration of myosin; *P* = 1 + [m]\**K*_W_(1 + *K_S_*); and *Q* = 1 + [m]\**K*_W_. *K*_B_ at high calcium and *K*_S_ were set to 20 and 18, respectively, based on (4). *nH*, *K*_W_, and *K*_T_ were determined in the presence and absence of calcium by fitting each titration curve independently.

### Modified fitting approach

To improve the resolution of the parameters extracted from fitting of the data, we modified the classic McKillop and Geeves approach:

1. We performed an additional titration at an intermediate calcium concentration (pCa 6.25).
2. The data for curves collected at three different calcium concentrations were globally fit using least squares optimization, and the parameters that minimized the aggregate error in all three data sets were determined. In global fitting, individual parameters can be shared between data sets, reducing the number of free variables and increasing the power to precisely measure parameter values. Here, *K*_W_ and *nH* were shared parameters among all three curves; however, this approach does not require sharing these parameters.
3. We used an annealing routine in our fitting procedure to avoid biasing the fit toward the initial guesses used in the fitting. The annealing routine ensures that the best-fit parameters are obtained from a global rather than local minimum.

### Hypothesis testing and statistics

One limitation of the original formulation of the three-state model is that it lacks procedures for calculating uncertainties and statistical hypothesis testing. Here, we used the well-established technique of bootstrapping simulations to calculate 95% confidence intervals (12, 13). In the bootstrapping method, the original data set is randomly resampled to generate synthetic data sets, each containing the same number of points as the original data set. Each synthetic data set is then fit to determine the best-fit values of the parameters for that data set. 95% confidence intervals are defined as the interval over which 95% of the simulated parameter values are found. Note that the number of bootstrapping simulations required will depend on the noise in the data as well as the number of sampled points. As such, one should empirically determine the number of simulations required for stable convergence of the confidence intervals. We have found that for the experiments described here, 1000 rounds of bootstrapping simulations were sufficient (Fig. 2A).

**Figure 2.**
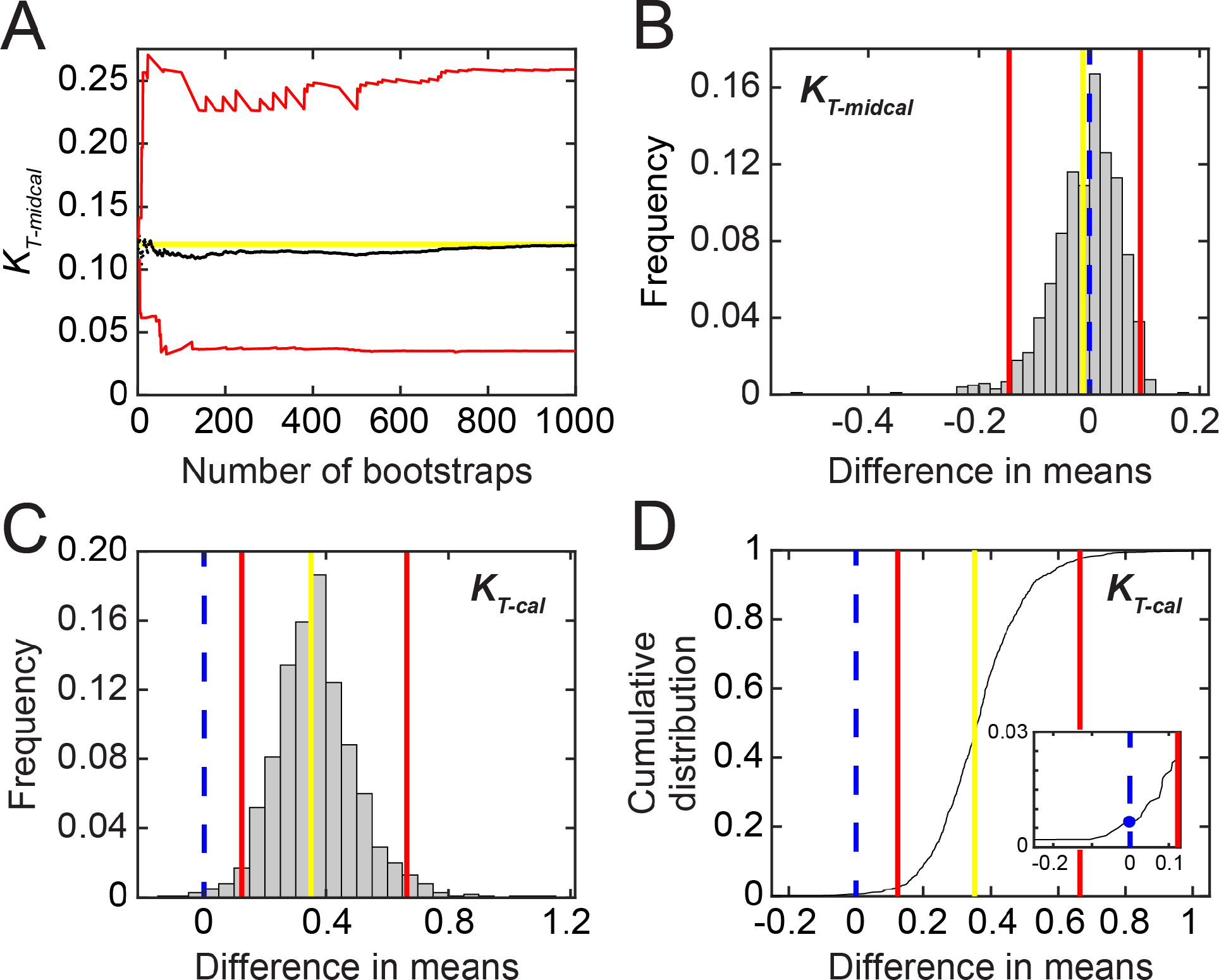
Hypothesis testing. **A.** Values of the average (black) and upper and lower bounds (red) of the 95% confidence interval for *K*_*T-midcal*_ as a function of the number of bootstraps performed during error estimation. The yellow line marks the measured best-fit value. **B-C.**Histograms of the test statistic (i.e., the difference in means between parameters determined for WT and ΔE160) for *K*_*T-midcal*_ (B) and *K*_*T-cal*_ (C). Vertical lines represent the measured difference in means (yellow), the bounds of the 95% confidence interval (red), and the null hypothesis (blue dashed). The difference in parameter values is statistically significant at the 95% confidence level if the null hypothesis falls outside the 95% confidence interval, as in C. **D.** Cumulative distribution of the difference in means between WT and ΔE160 calculated from the bootstrapping simulations for *K*_*T-cal*_. Vertical lines are the same as in C. Inset highlights the determination of the p-value (blue) from the value of the cumulative distribution at x = 0. This value is multiplied by two to make the test equivalent to a two-tailed test.

For statistical hypothesis testing, we defined a test statistic as the difference between parameter values in the perturbed and unperturbed systems (14). For example, to determine whether there is a statistically significant difference in the value of *K*_W_ for a mutant protein relative to wild-type (WT), the test statistic is H = *K*_W_(WT) – *K*_W_(mutant). The value of the test statistic is calculated for the real data and for pairs of *K*_W_ values drawn from the bootstrapping simulations of the data. The interval within which 95% of the test statistic values from the bootstrapped simulations fall is defined as the 95% confidence interval. If the null hypothesis (i.e., H_0_=0) is not contained within the 95% confidence interval, then the null hypothesis is rejected, and *K*_W_(WT) ≠ *K*_W_(mutant) with a p-value < 0.05 (Fig. 2B-C). The p-value can be calculated from a cumulative distribution of the test statistic by finding the largest interval that does not contain the null hypothesis (Fig. 2D). This value is multiplied by two to make the test equivalent to a two-tailed test.

## Results and Discussion

Here we present a methodology and MATLAB-based computational tool for analyzing biochemical measurements of thin-filament positioning. This procedure builds on the formalism developed by McKillop and Geeves (4), extending it to allow for improved precision of parameter values, estimations of uncertainties, and statistical hypothesis testing. The basic workflow for the analysis follows:

1. Measure the value of the equilibrium constant between the blocked and closed states, *K_B_*, using a stopped-flow kinetic technique (see Supporting Materials).
2. Perform steady-state fluorescence titrations measuring myosin binding to regulated thin filaments (RTFs; see Supporting Materials). The titrations should be carried out at three separate calcium concentrations (i.e., low, high, and intermediate calcium concentrations).
3. Normalize the data from each technical replicate before pooling the data.
4. Globally fit the pooled titration data set to determine the best fit parameters and calculate confidence intervals from bootstrapping simulations.
5. For a given perturbation, statistically test for differences between individual parameters obtained from the fitting.

A detailed user guide describing how to perform each of these steps, along with all of the data used to generate the figures in this manuscript is provided with the computational tool.

### Results obtained using the traditional fitting procedure and the estimation of uncertainties

The individual equilibrium constants that define the positioning of cardiac tropomyosin along the thin filament were determined using tissue-purified porcine cardiac myosin and actin and recombinant human troponin and tropomyosin (see Supporting Materials). The equilibrium constant for the transition between the blocked and closed states, *K_B_*, was determined by performing stopped-flow kinetic measurements. We measured *K*_*B*_ = 0.3 ± 0.2, in agreement with the previously determined value (4, 15). Although there is some variance between technical replicates for *K*_B_, the values of parameters obtained from fitting of the titration data are relatively insensitive to small changes in the value of *K_B_*.

The equilibrium constants for the transition between the closed and open states, *K_T_*, and for myosin weak binding, *K_W_*, were determined by performing titrations of RTFs with myosin at high (pCa 3) and low (2 mM EGTA) calcium. The titration curves were fit independently by Equation 1 (4). To ensure that the fitting was not biased by the initial guesses of parameter values, we modified the fitting procedure described in (4) to incorporate an annealing routine that iteratively fits the data with altered initial guesses. The values obtained from these fits are shown in Figure 3A, and they are consistent with values obtained previously with skeletal and cardiac muscle proteins (15–17). The original formulation of the McKillop and Geeves model did not include a methodology for determining uncertainty in fitted parameters. We developed a procedure to calculate 95% confidence intervals via bootstrapping of the original data set (12, 13). With this procedure, we saw that the classic approach gave considerable uncertainty in the values of the fitted parameters. For example, for the data collected at low calcium, we obtained uncertainties much larger than the measured parameters for both *K*_*W*_ (0.13 (−0.03/ +1.26)) and *nH* (6 (−2/+6)).

**Figure 3.**
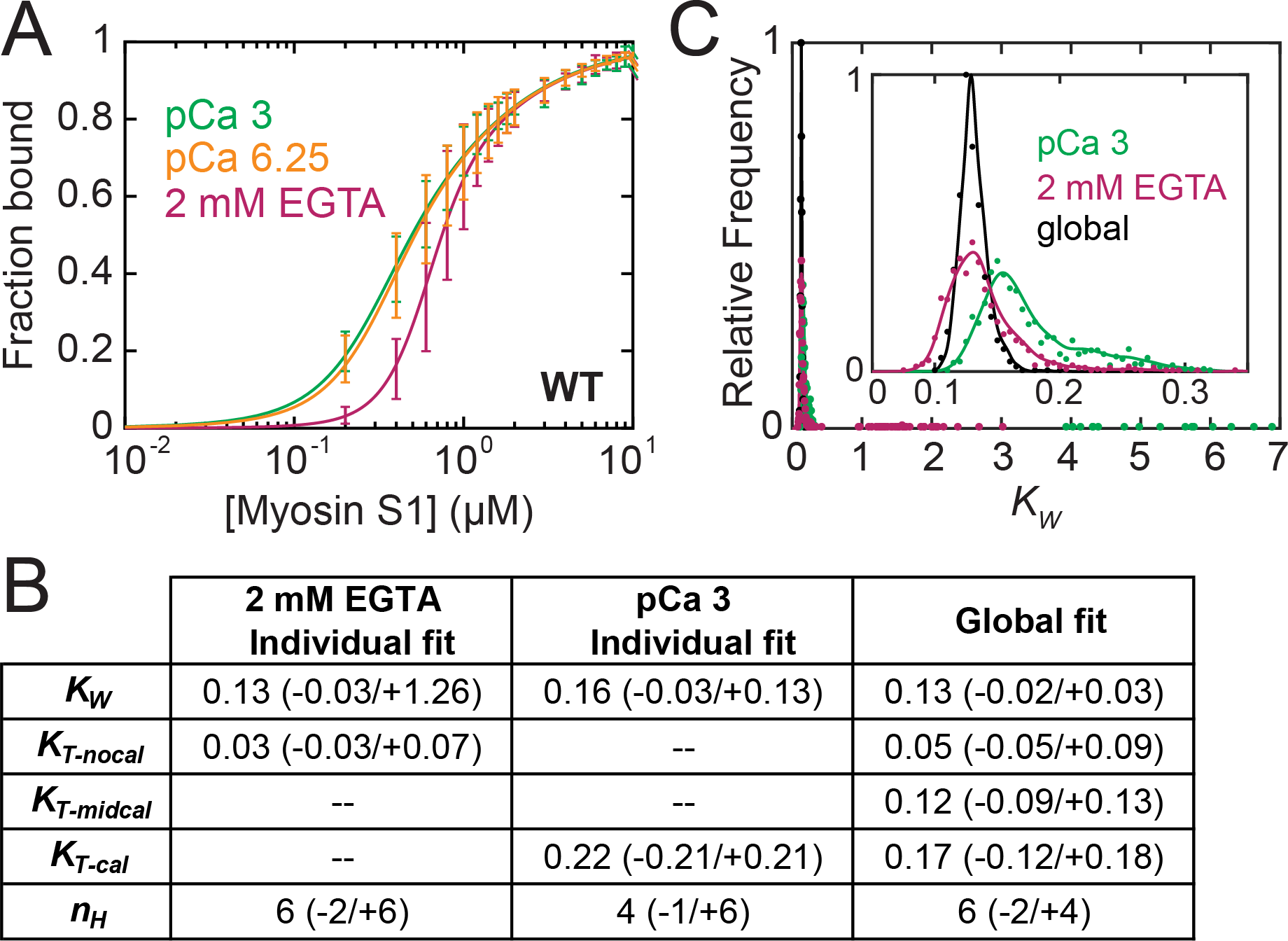
Fitting the three-state model to fluorescence titration data. **A.** Steady-state titrations of regulated thin filaments with myosin for the WT protein conducted at three distinct calcium concentrations: saturating calcium (pCa 3, green), intermediate calcium (pCa 6.25, orange), and no calcium (2 mM EGTA, magenta). Curves are fits of the data. Error bars show the standard deviation of five technical replicates. **B.** Table of parameter values obtained from individual fits of data collected at 2 mM EGTA or pCa 3 compared to those obtained from our global fitting method, which includes pCa 6.25. Values in parentheses indicate the 95% confidence intervals. **C.** Histograms showing the frequency of *K*_*W*_ values determined from 1000 bootstrapping simulations. Points are the values from the global fitting (black) and from the individual fits of data collected at 2 mM EGTA (magenta) and pCa 3 (green). Lines are inserted to guide the eye. The inset shows the distributions near the mean values of *K*_*W*_. These data demonstrate that global fitting reduces the uncertainty in the measurement.

### Global fitting improves the resolution of fitted parameters

To improve the resolution of parameters derived from the fitting of the data, we adopted two modifications from the traditional approach. First, we performed an additional titration at an intermediate calcium concentration (pCa 6.25). The inclusion of an additional titration curve at an intermediate calcium is advantageous because some perturbations shift the calcium sensitivity at intermediate calcium concentrations but not under fully activating or deactivating conditions (18). Second, titration curves collected at the three different calcium concentrations were fit globally rather than individually. In the global fitting, the values of *K_W_*, *K_S_*, and *nH* were shared between the fits of all curves. Global fitting of the three curves (Fig. 3A) yielded tighter 95% confidence intervals for both *K*_*W*_ and *nH* (Fig. 3). The confidence interval for *K*_*W*_ was 0.13 (−0.02/+0.03), compared to 0.13 (−0.03/+1.26) for the individually fit curve at low calcium. Similarly, the confidence interval for *nH* for the individually fit data was 6 (−2/+6), while the confidence interval for the globally fit data was 6 (−2/+4). These data demonstrate the improved resolution of this approach.

### Hypothesis testing and its application to determining the biochemical mechanism of a mutation that causes hypertrophic cardiomyopathy

To demonstrate the utility of our approach for resolving the effects of molecular-based changes in cardiac thin-filament regulation, we examined a point mutation that causes familial hypertrophic cardiomyopathy (HCM), ΔE160 in troponin T (11). Clinical studies have shown that this mutation causes pronounced ventricular hypertrophy and sudden cardiac death, with half of the patients not surviving past age 40 (11). Previous studies of the ΔE160 mutation in muscle fibers (19, 20), transfected myotubes (21), and purified proteins (22) have shown that the ΔE160 mutation causes increased activation of contractility. However, the biochemical mechanism of this activation is not well understood.

We applied our methodology to determine which transitions involved in thin-filament regulation (Fig. 1) are affected by the ΔE160 mutation. The goal of this analysis is not an exhaustive characterization of the mutant; rather, it is to demonstrate the utility of this approach for statistically assessing the effects of perturbations on thin-filament regulation. From stopped-flow kinetic measurements (Fig. 4A), we obtained a *K*_*B*_ of 0.2 ± 0.1 for ΔE160, which is indistinguishable from the value obtained for wild-type (WT) troponin (0.3 ± 0.2; p=0.41). We also performed fluorescence titrations (Fig. 4B) to measure steady-state actomyosin binding and determine *K*_*W*_, *K_T_*, and *nH*. We used the computational tool to calculate parameter values and their corresponding uncertainties from global fitting of the titration curves (Fig. 4C).

**Figure 4.**
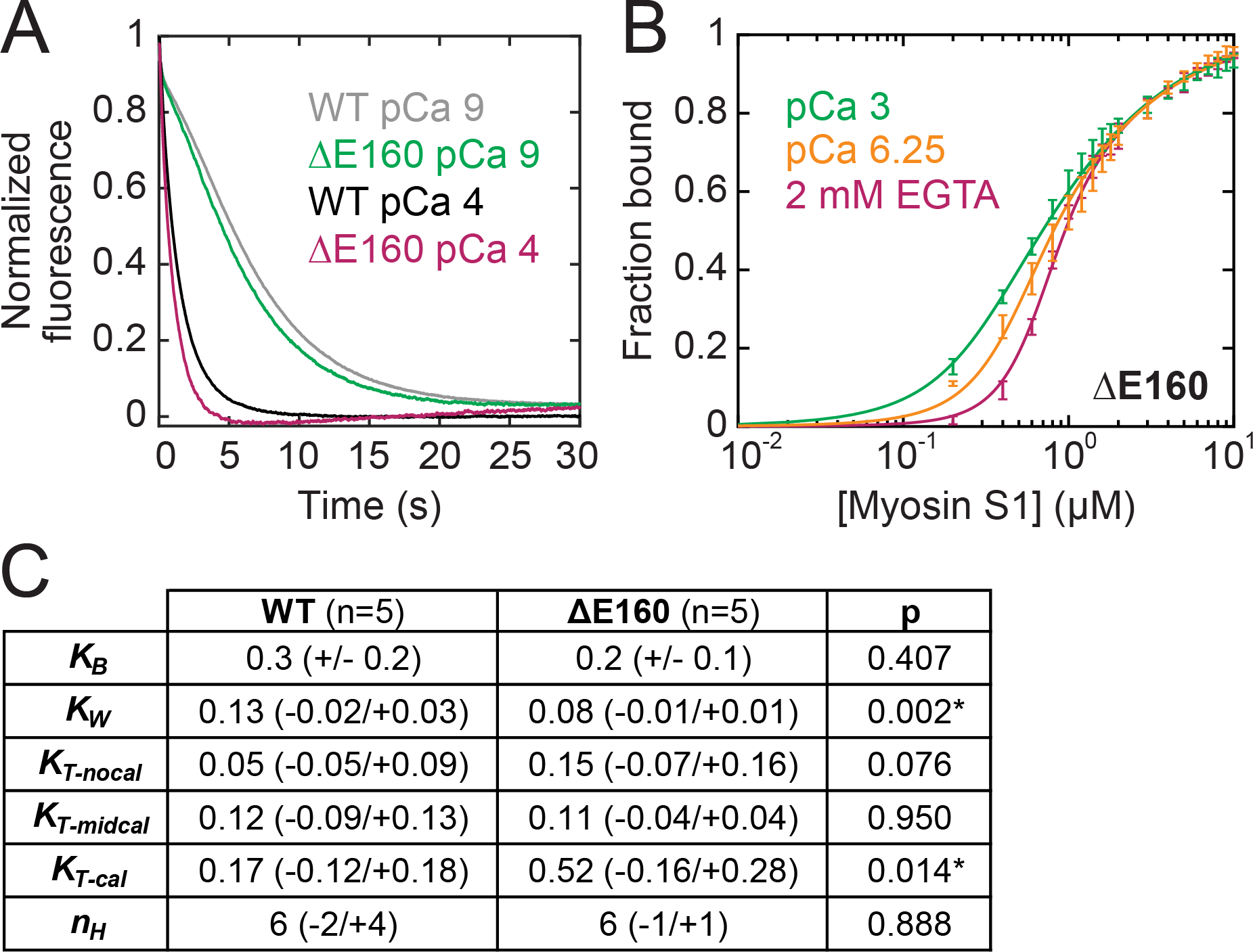
Effects of ΔE160 mutation on thin-filament regulation. **A.** Normalized stopped-flow fluorescence traces of regulated thin filaments binding to myosin. The pyrene fluorescence is quenched at a higher rate at saturating calcium (pCa 4, magenta/black) than at low calcium (pCa 9, green/gray). The traces for WT (black/gray; average of n=6 curves each) and the ΔE160 mutant (magenta/green; average of n=4 curves each) are similar at each calcium concentration. **B.** Steady-state titrations of regulated thin filaments with myosin for the mutant protein conducted at three distinct calcium concentrations: saturating calcium (pCa 3, green), intermediate calcium (pCa 6.25, orange), and low calcium (2 mM EGTA, magenta). Curves are fits of the data. Error bars show the standard deviation of five technical replicates. **C.** Table of parameter values obtained for WT and ΔE160 troponin complexes from stopped-flow measurements (for *K_B_*) and using the computational tool (for all others). Values in parentheses indicate the 95% confidence intervals. Asterisks indicate statistical significance at the 95% confidence level.

To determine whether there are statistically significant differences between the parameters for WT troponin and the ΔE160 mutant, we applied the hypothesis testing methodology described in the Materials and Methods. We found that the mutant showed a threefold increase in *K*_*T*_ at high calcium (0.17 (−0.12/+0.18) for WT versus 0.52 (−0.16/+0.28) for ΔE160; p=0.014). This threefold increase in *K*_T_ would result in an increased population of force-generating cross bridges at high calcium. We also found a slight, but statistically significant, decrease in *K*_*W*_ (0.13 (−0.02/+0.03) for WT versus 0.08 (−0.01/+0.01) for ΔE160; p=0.002). We saw that at low calcium, *K*_*T*_ was 3-fold larger for the mutant (p=0.076); however, this difference was not significant at the 95% confidence level. We did not detect statistically significant differences in the values of *nH* (p=0.89), *K*_*B*_ (p=0.41), or *K*_*T-midcal*_ (p=0.95). The net effect of these changes would be to increase activation, which is consistent with the hypercontractility associated with HCM. Taken together, these data demonstrate the power of this approach for hypothesis testing and for determining the biochemical mechanism of perturbations of thin-filament regulation.

### Application of the hypothesis testing and uncertainty estimation to other systems

The computational tool for hypothesis testing and confidence interval estimation from bootstrapping simulations is not limited to analysis of fluorescence titrations, but it can be broadly applied to other data sets as well. We have supplied a stand-alone version of this section of the code for examining the mean and median values (i.e., data frequently used for single cell measurements) so that others can apply it to their experimental system. This methodology is useful for data sets for which the form of the underlying distribution is either unknown or not normal, such as single-molecule data (23) and single-cell studies, as demonstrated in (18).

### Conclusion

Here, we have demonstrated a method for extending the utility of the McKillop and Geeves (4) approach to understanding thin-filament regulation, and we have provided a well-documented, accessible computational tool to implement this methodology. Our approach extends the McKillop and Geeves approach to include a method for calculating confidence intervals and performing statistical tests. This methodology allowed us to resolve the molecular effects of a mutation that causes hypertrophic cardiomyopathy. This tool should be useful for studying physiological and pathological changes in muscle, as well as for testing new therapies that target muscle regulation.

## Code Availability

The computational tool can be downloaded from GitHub at: https://github.com/GreenbergLab/Thin_Filament_Fitting

We also provide an in-depth user guide along with the raw data used in the examples presented here.

## Acknowledgements

Funding for this project was provided by the National Institutes of Health (R00HL123623, R01HL141086 to M.J.G., T32EB018266 to S.R.C.), and the March of Dimes Foundation (FY18-BOC-430198 to M.J.G.). We thank Tommy Blackwell and Tom Stump for critical evaluation of the code.

## Author contributions

S.K.B. performed and analyzed the fluorescence experiments and developed code. S.R.C. helped with the early development and testing of the MATLAB code. L.G. generated mutant protein. M.J.G. oversaw the project and developed the early code for analyzing data. S.K.B. and M.J.G. wrote the first draft of the paper, and all authors contributed to the final draft.

