## Supporting Materials for "Computational tool to study perturbations in muscle regulation and its application to heart disease"

### Protein expression and purification

Cardiac ventricular myosin and actin were prepared from cryoground porcine ventricles (Pelfreez) as previously described (1). Cardiac actin was labeled with N-(1-pyrenyl)iodoacetamide (pyrene-actin) as described in (2, 3). The concentration of pyrene-actin was determined by measuring the absorbances at 290 nm and 344 nm. Myosin subfragment-1 (S1) was prepared from full-length myosin by chymotrypsin digestion (4), and its purity was assessed using SDS-PAGE.

Recombinant human tropomyosin was expressed in BL21-CodonPlus cells (Agilent) and purified using established protocols (5). The concentration of tropomyosin was determined by measuring the absorbance at 280 nm.

Human recombinant troponin I, troponin C, and troponin T were expressed in BL21-CodonPlus cells (Agilent), complexed, and purified using established protocols (6), with the modification that the final complexed protein was purified using a MonoQ column. Troponin complex concentration was measured spectroscopically using the Bradford reagent (Coomassie Plus Assay kit, Thermo Scientific). The ΔE160 mutation in troponin T was introduced using the QuikChange Site-Directed Mutagenesis Kit (Agilent) and verified by sequencing (GENEWIZ). The ΔE160 protein was expressed, purified, and complexed into functional troponin units using the same procedure as the WT protein.

### Stopped-flow kinetic measurements of $K_B$

Stopped-flow transient kinetic measurements were carried out in buffer containing 60 mM MOPS, 200 mM KCl, 5 mM $MgCl_2$, 1 mM DTT, 2 mM EGTA, and either 5.2 μM

$CaCl_2$ (pCa 9) or 2.15 mM $CaCl_2$ (pCa 4). Total concentrations of calcium, magnesium, ATP, and EGTA added were calculated using MaxChelator (7). Prior to measurements, tropomyosin was reduced by addition of 50 mM DTT, heated at 56 °C for 5 min, and allowed to cool to room temperature before centrifuging 30 min at 436000 x g at 4 °C in an Optima MAX-TL Ultracentrifuge (Beckman Coulter) equipped with a TLA 120.2 rotor (Beckman Coulter). Phalloidin-stabilized, pyrene-labeled actin filaments were prepared by incubating 50 μM pyrene-actin with 55 μM phalloidin in low-calcium buffer at least 10 min at room temperature.

$K_B$, the equilibrium constant between the blocked and closed states, was calculated by measuring the rates of myosin strong binding to pyrene-labeled regulated thin filaments (RTFs) in the presence and absence of calcium, as previously described (8). Pyrene was excited at 365 nm, and the fluorescence emission was detected using a 395-nm cutoff filter. Fluorescent RTFs (2.5 μM phalloidin-stabilized pyrene-actin and 1 μM troponin and tropomyosin) were rapidly mixed with 0.25 μM myosin subfragment-1 (S1) using an SX20 stopped-flow mixer (Applied Photophysics). Apyrase (0.02 units/mL) was added to each protein solution before mixing to ensure the absence of nucleotide (i.e., ATP and ADP). All concentrations refer to the final concentrations after mixing.

When myosin S1 was rapidly mixed with RTFs, the pyrene fluorescence in the thin filaments was quenched by myosin strong binding, yielding an exponential decrease in fluorescence. Rate constants, $k$, for actomyosin binding were obtained by fitting a single-exponential curve to the average of at least three kinetic traces for each calcium concentration. $K_B$ was calculated by taking the ratio of the rate constants at low (pCa 9) and high (pCa 4) calcium (8):

$$\frac{k(-Ca^{2+})}{k(+Ca^{2+})} = \frac{K_B}{1+K_B} \quad \text{Equation S1}$$

The reported $K_B$ values are the average of at least four independent experiments, and reported errors are the standard deviations between these values. Statistical significance between $K_B$ values was evaluated using a Student's two-tailed t-test.

**Measurement of steady-state myosin binding to regulated thin filaments to determine $K_T$, $K_W$, and $n$**

The parameters $K_T$, $K_W$, and $nH$ were calculated from equilibrium titrations measuring the concentration-dependent binding of myosin to pyrene-labeled RTFs (8). Myosin strong binding to the pyrene-labeled RTFs quenches the fluorescence, thus the fractional change in fluorescence was measured as a function of myosin added at three calcium concentrations. Fluorescence titrations were performed in buffer containing 60 mM MOPS, 200 mM KCl, 5 mM $MgCl_2$, 1 mM DTT, and either 2 mM EGTA (low $Ca^{2+}$), 2 mM EGTA and 1.19 mM $CaCl_2$ (pCa 6.25), or 1 mM $CaCl_2$ (pCa 3). Reduced tropomyosin and phalloidin-stabilized pyrene-actin were each prepared as described above. RTFs (0.5 μM phalloidin-stabilized pyrene-actin and 0.27 μM troponin and tropomyosin) were continuously stirred and myosin S1 was added in the presence of 2 mM ADP, as well as 50 μM $P^1$,$P^5$-di(adenosine-5′) pentaphosphate (Ap5a), 1 μM hexokinase, and 2 mM glucose to maintain the nucleotide in the ADP state. Myosin S1 was added at 1-minute intervals in increments of 0.2 μM up to 2 μM, then at 1-μM increments up to 10 μM. The steady-state pyrene fluorescence was measured at 1-minute intervals. Five technical replicates per condition were collected.
